# SARS-CoV-2 Infection And Longitudinal Fecal Screening In Malayan Tigers (*Panthera tigris jacksoni*), Amur Tigers (*Panthera tigris altaica*), And African Lions (*Panthera leo krugeri*) At The Bronx Zoo, New York, USA

**DOI:** 10.1101/2020.08.14.250928

**Authors:** Susan L. Bartlett, Diego G. Diel, Leyi Wang, Stephanie Zec, Melissa Laverack, Mathias Martins, Leonardo Cardia Caserta, Mary Lea Killian, Karen Terio, Colleen Olmstead, Martha A. Delaney, Tracy Stokol, Marina Ivančić, Melinda Jenkins-Moore, Karen Ingerman, Taryn Teegan, Colleen McCann, Patrick Thomas, Denise McAloose, John M. Sykes, Paul P. Calle

**Author notes:** ZooRadOne, Plainfield, IL 60544, USA. These authors contributed equally to this work (Susan L. Bartlett, Diego G. Diel, Leyi Wang). Correspondence should be directed to Dr. Bartlett.

## Abstract

Severe Acute Respiratory Syndrome Coronavirus-2 (SARS-CoV-2) emerged as the cause of a global pandemic in 2019-2020. In March 2020 New York City became the USA epicenter for the pandemic. On March 27, 2020 a Malayan tiger (*Panthera tigris jacksoni*) at the Bronx Zoo in New York City developed a cough and wheezing with subsequent inappetence. Over the next week, an additional Malayan tiger and two Amur tigers (*P. t. altaica*) in the same building and three lions (*Panthera leo krugeri*) in a separate building also became ill. The index case was immobilized, and physical examination and bloodwork results were unremarkable. Thoracic radiography and ultrasonography revealed peribronchial cuffing with bronchiectasis, and mild lung consolidation with alveolar-interstitial syndrome, respectively. SARS-CoV-2 RNA was identified by real-time, reverse transcriptase PCR (rRT-PCR) on oropharyngeal and nasal swabs and tracheal wash fluid. Cytologic examination of tracheal wash fluid revealed necrosis, and viral RNA was detected in necrotic cells by *in situ* hybridization, confirming virus-associated tissue damage. SARS-CoV-2 was isolated from the tracheal wash fluid of the index case, as well as the feces from one Amur tiger and one lion. Fecal viral RNA shedding was confirmed in all seven clinical cases and an asymptomatic Amur tiger. Respiratory signs abated within 1-5 days for most animals, though persisted intermittently for 16 days in the index case. Fecal RNA shedding persisted for as long as 35 days beyond cessation of respiratory signs. This case series describes the clinical presentation, diagnostic evaluation, and management of tigers and lions infected with SARS-CoV-2, and describes the duration of viral RNA fecal shedding in these cases. This report documents the first known natural transmission of SARS-CoV-2 from humans to animals in the USA, and is the first report of SARS-CoV-2 in non-domestic felids.

## INTRODUCTION

In December 2019 cases of pneumonia of unknown etiology occurred in people in Wuhan, China.^30^ By January 2020 the cause of the infection was identified as a novel coronavirus named Severe Acute Respiratory Syndrome Coronavirus 2 (SARS-CoV-2), and the resulting disease referred to as COVID-19.^6^ The emergence of this virus was associated with Wuhan’s Huanan Seafood Wholesale Market, which also sold various species of live wild animals.^2,33^ The virus shares more than 96% homology with a coronavirus isolated from bats (BatCoV RaTG13).^8,16,36^ The exact transmission route from bats to people is unknown, though transmission through one or more intermediate hosts is suspected. After rapid global spread, the outbreak was declared a pandemic by the World Health Organization on March 11, 2020.^31^ SARS-CoV-2 was first documented in people in the United States of America (USA) in late January 2020 in Washington State, with the first confirmed case in New York State on February 29, 2020 in New York City (NYC).^4^ New York State subsequently became the epicenter of the pandemic in the USA, with over 400,000 human infections across the state as of July 13, 2020, approximately half of which occurred in NYC.^13^ This case series describes detailed clinical findings, outcomes, and patterns and duration of fecal viral shedding in relation to cessation of clinical signs due to infection with SARS-CoV-2 in Malayan (*Panthera tigris jacksoni*) and Amur (*P. t. altaica*) tigers and African lions (*Panthera leo krugeri*) at the Wildlife Conservation Society’s (WCS) Bronx Zoo in New York City, New York, USA.

## CASE SERIES

The WCS operates four zoos and an aquarium in New York City, all accredited by the Association of Zoos and Aquariums (AZA). The Bronx Zoo houses snow leopard (*Panthera uncia*), cheetah (*Acinonyx jubatus*), clouded leopard (*Neofelis nebulosa*), Amur leopard (*Panthera pardus orientalis*), puma (*Puma concolor*), serval (*Leptailurus serval*), Malayan and Amur tigers, and lions. The Central Park Zoo houses snow leopard; the Queens Zoo houses Canada lynx (*Lynx canadensis*); the Prospect Park Zoo houses black-footed cat (*Felis nigripes*) and Pallas cat (*Otocolobus manul*). The Bronx Zoo exhibits tigers in two locations, Tiger Mountain and Wild Asia, which are approximately 3,000 feet apart. Two Malayan tigers [T1 and T2] and three Amur tigers [T3-T5] are housed individually at Tiger Mountain and have no direct contact with each other, but are rotated through shared enclosures, outdoor holding yards, and exhibits (Fig. 1). One Malayan and two Amur tigers are housed in Wild Asia. The Bronx Zoo also houses three African lions (L1-L3) in the African Plains exhibit, which is located 1,500 feet from Tiger Mountain, and 2,300 feet from Wild Asia. The lions are housed individually overnight, but exhibited in pairs during the day. L2 is exhibited with L1 or L3 on an alternating basis. L1 and L3 are never in direct contact (Fig. 1). The tigers and lions are all adults, ranging in age from 4-15 years old and were born at the Bronx Zoo, with the exception of T3 who arrived in 2015.

**Figure 1.**
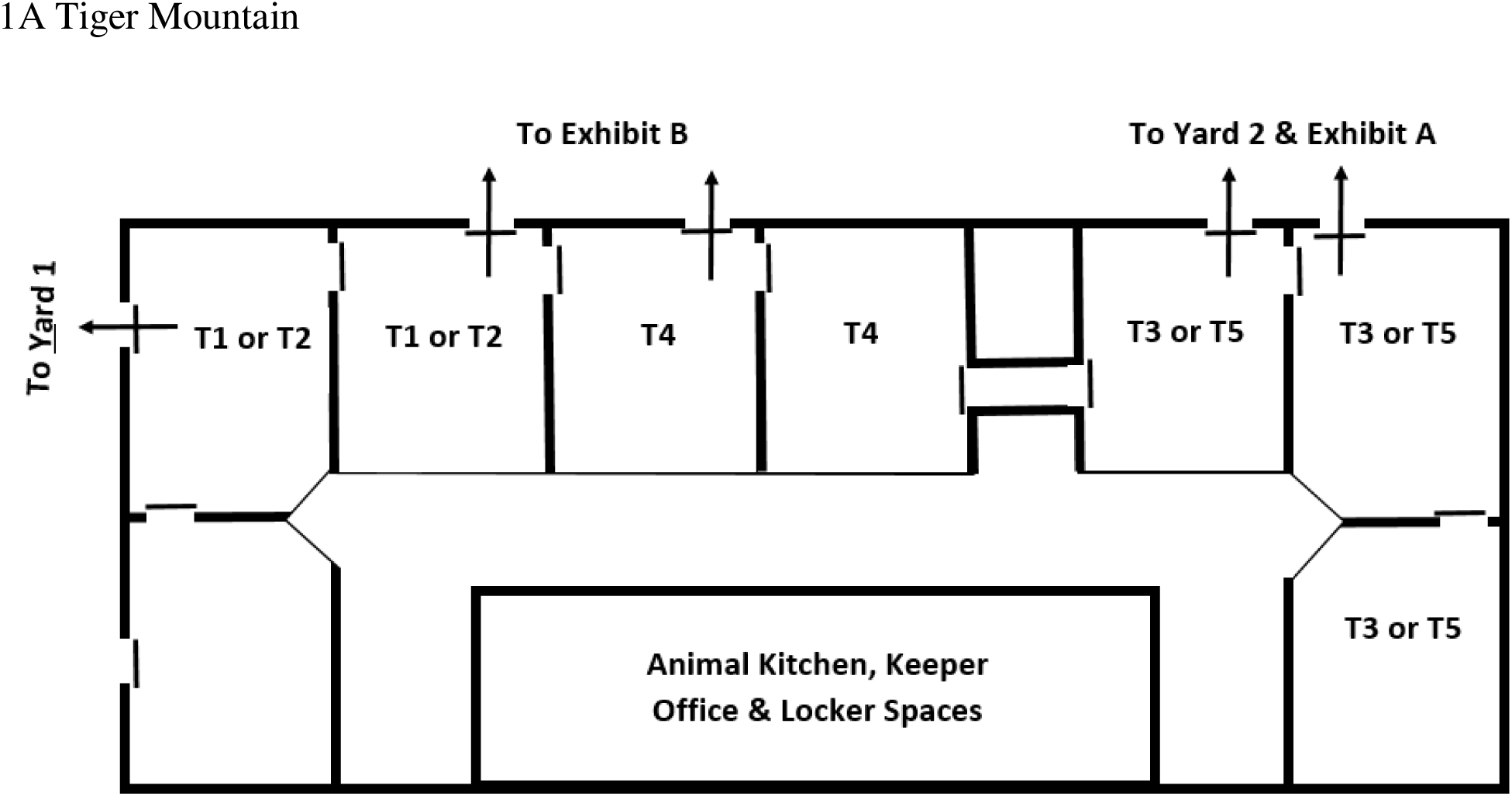
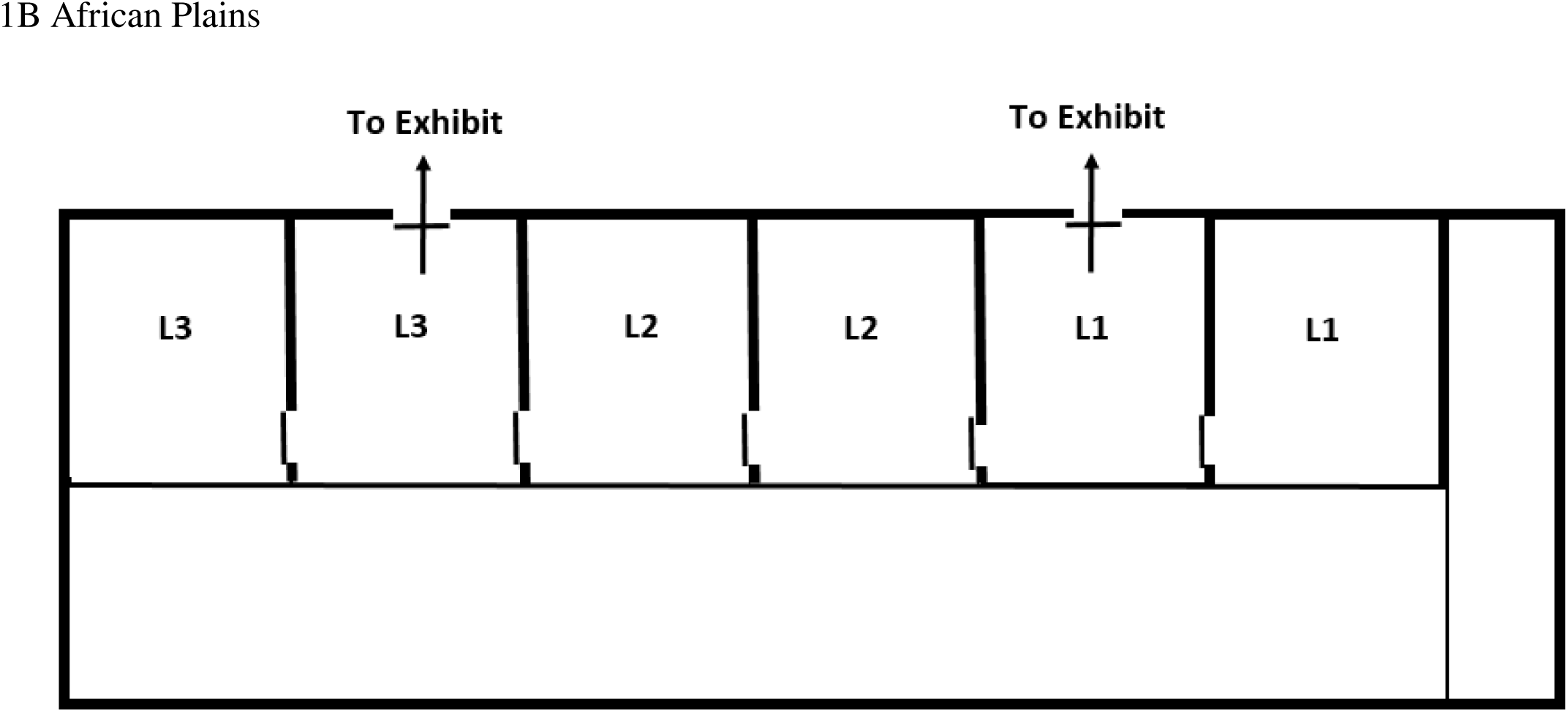
Schematic diagram of Tiger Mountain (A) and African Plains (B) facilities at the Bronx Zoo where tigers (T1-T5) were housed and exhibited individually, and lions (L1-L3) were housed individually and exhibited in alternating pairs (L1/L2 or L2/L3), respectively.

On March 27, 2020 (Day 0), a 4-yr-old female Malayan tiger (T1) from Tiger Mountain developed a cough, which varied from dry to wet, with occasional wheezing. The episodes of coughing lasted 20-30 seconds and were heard intermittently throughout the day. The clinical signs persisted the following day, so treatment was initiated with amoxicillin/clavulanic acid (375 mg tablets, Zoetis Inc., Kalamazoo, Michigan 49007, USA; 12.8 mg/kg PO, BID for 21 days). In the subsequent week, the second Malayan tiger (T2) and two Amur tigers (T3 and T4) in Tiger Mountain, and the three lions in African Plains, all developed similar clinical signs (Table 1). No clinical signs were observed in one Amur tiger (T5) in Tiger Mountain or any of the tigers in Wild Asia. All affected animals were treated with amoxicillin/clavulanic acid (11.5 – 14 mg/kg PO BID for 14 days) at the onset of clinical signs. They remained eupneic with no ocular or nasal discharge throughout their illness, and their behavior was otherwise normal. Two tigers (T2 and T5) each had one mild episode of unilateral epistaxis, occurring on March 26 (Day -1) and April 2 (Day 6) respectively. The index case (T1) had an increased frequency of coughing by Day 4, with bouts commonly following periods of increased activity, and a decreased appetite.

**Table 1.**
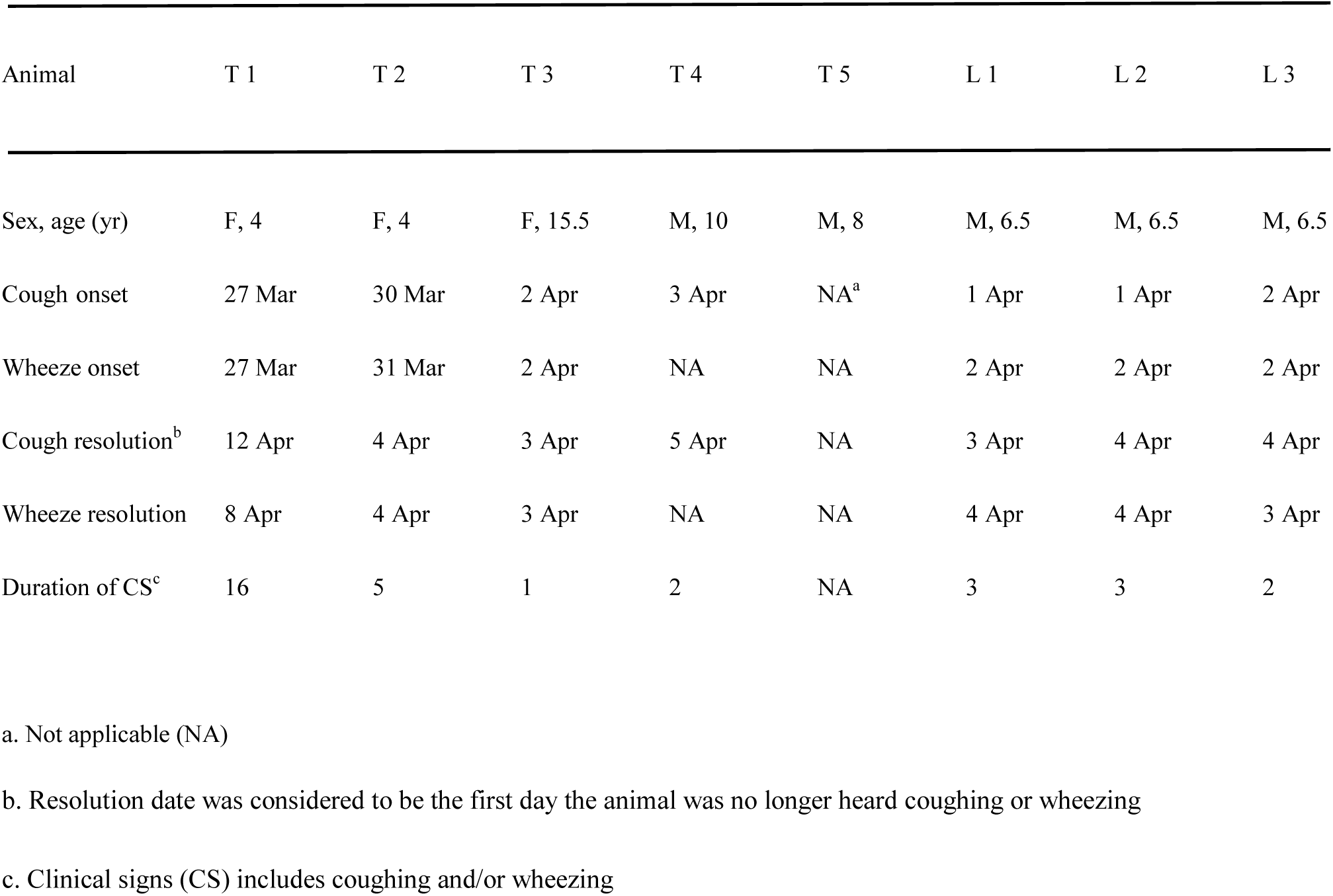
Signalment, onset, and duration of respiratory signs in SARS-CoV-2 infection of Malayan (*Panthera tigris jacksoni*) and Amur (P. *t. altaica*) tigers and African lions (*Panthera leo krugeri*). Note Tiger 5 never developed clinical signs.

Due to the persistence of clinical signs, T1 was immobilized for treatment and diagnostic evaluation on Day 6. Personal protective equipment (PPE) for staff members (veterinarians, technicians, and animal care staff) working around the tiger’s head included N95 masks, full face shields and disposable examination gloves; surgical masks and examination gloves were worn by all others participating in the procedure. The animal was anesthetized by dart injection of medetomidine (ZooPharm, Laramie, Wyoming 82070, USA; 0.03 mg/kg IM) and ketamine (ZooPharm; 3 mg/kg IM), followed by nasal flow-by isoflurane (MWI, Boise, Idaho 83705, USA; 5%) and intravenous diazepam (Hospira, Inc., Lake Forest, Illinois 60045, USA; 0.057 mg/kg IV) after light anesthesia was achieved. Rectal temperature obtained immediately after induction was normal (101.4 °F, normal range 99.3 – 102.9 °F), ^7^ and auscultation of the heart and lungs was unremarkable. A tracheal wash and sample collection was then performed as follows: Lidocaine (MWI; 40 mg topically) was applied to the laryngeal folds to minimize a cough reflex. A sterile 20 French red rubber tube (Amsino International, Inc., Pomona, California 91768, USA) was inserted into the proximal trachea, using sterile technique. Then, 60 ml of sterile saline (Hospira) was instilled into the trachea; approximately half of the instilled fluid was retrieved on aspiration. The fluid appeared flocculent and pink-tinged. Endotracheal intubation with a 16mm internal diameter tube was then performed and isoflurane administered (2.5-5%) for maintenance of anesthesia. Physical examination, thoracic and abdominal radiography and ultrasonography were performed. Blood samples were taken for clinical pathologic testing, and duplicate oropharyngeal and nasal samples were collected using polypropylene swabs (Becton, Dickinson and Company, Sparks, Maryland 21152, USA) and placed into cryovials. While under anesthesia, the tiger was treated supportively with cefovecin sodium (Zoetis Inc.; 8 mg/kg SC), penicillin (Norbrook Laboratories Limited, BT34 Newry, Northern Ireland; 30,000 IU/kg SC), and lactated ringer’s solution (Dechra Veterinary Products, Overland Park, Kansas 66211, USA; 11.4 ml/kg SC). After administration of atipamezole (ZooPharm; 0.17 mg/kg IM), the animal recovered from anesthesia uneventfully. The entire procedure from anesthetic drug administration to arousal with control of the head was 97 min.

Physical examination revealed the animal was in good condition with no significant abnormalities. Opposite lateral and dorsoventral radiographs of the thorax demonstrated a generalized bronchial pattern with multifocal caudal peribronchiolar cuffing and bronchiectasis (Fig 2). Ultrasonographic examination of the left lung performed in right lateral recumbency revealed at least two small areas of consolidated peripheral lung and adjacent coalescent vertical B-lines, typical of alveolar-interstitial syndrome (AIS).^22^ Abdominal radiographic and ultrasonographic examinations were unremarkable. In-house blood smear evaluation revealed a normal estimated total white blood cell count (6.7 × 10^3^/µl; reference interval 6 – 14 × 10^3^/µl), but 25% of the lymphocytes showed reactive features (e.g. large in size with deeply basophilic cytoplasm).^25^ No other morphologic abnormalities were noted in cells in the smear. Results of a serum biochemical profile performed at a regional diagnostic laboratory (Antech, New Hyde Park, New York 11042, USA) were unremarkable.

**Figure 2.**
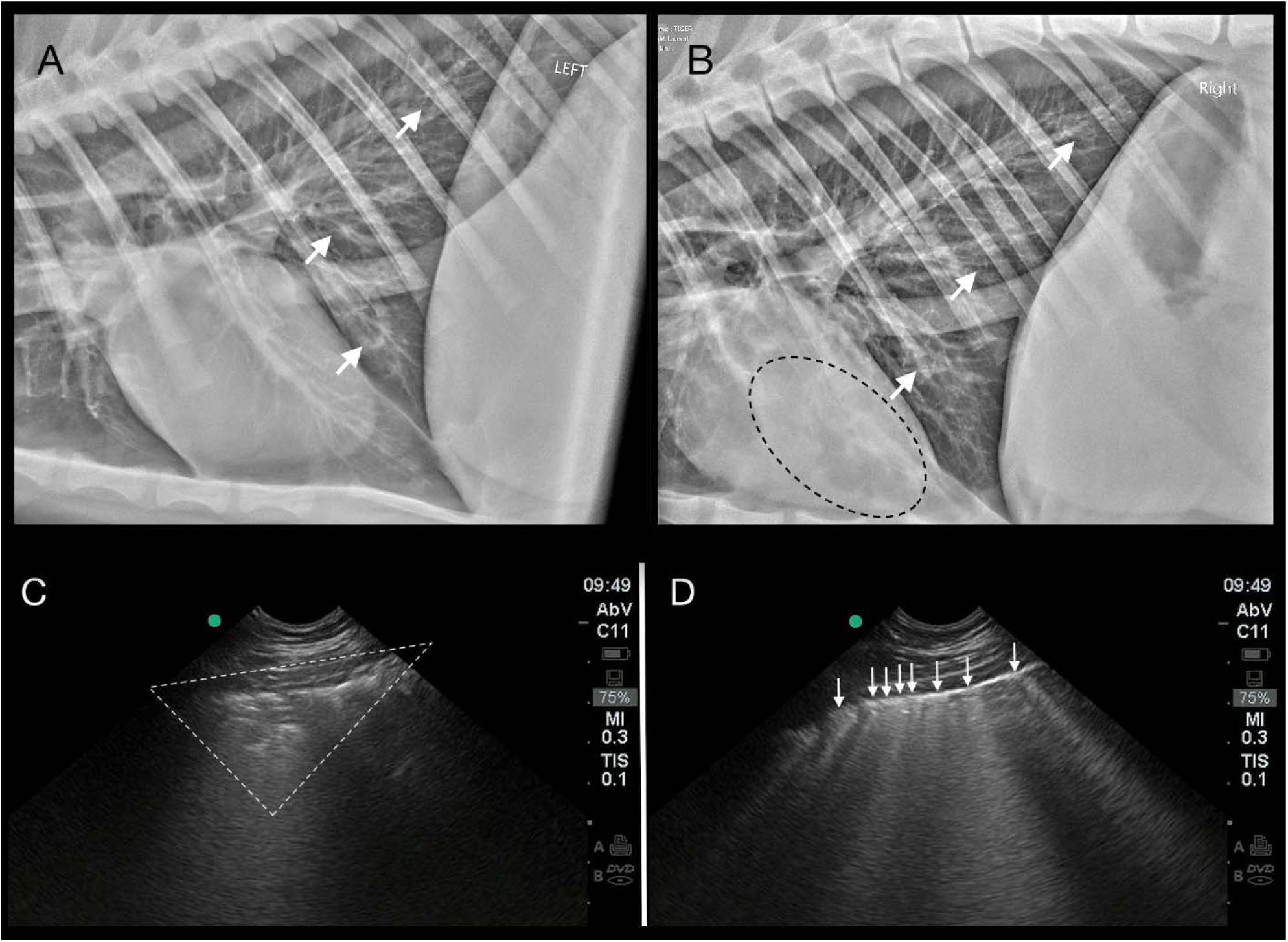
Thoracic imaging abnormalities in the index Tiger (T1) with SARS-CoV-2 infection. A generalized bronchial pattern with peribronchial cuffing and bronchiectasis (white arrows) is present in the caudal lung on left lateral (2A) and right lateral (2B) radiographs. Anesthesia-associated atelectasis is seen as an alveolar pattern superimposed over the heart (black dotted line) (2B). Pulmonary ultrasonography reveals peripheral consolidation (white dotted triangle) (2C), and coalescence of vertical B-lines (white arrows) (2D) indicating AIS (alveolar-interstitial syndrome).

T1 remained partially anorexic on Day 8, two days after the immobilization. The tiger was treated with maropitant (60 mg tablets; Zoetis Inc.; 1 mg/kg PO, SID for 7 days) and meloxicam (7.5 mg tablets; Zydus Pharmaceuticals (USA) Inc., Pennington, New Jersey 08534, USA; 0.2 mg/kg PO once, followed by 0.1 mg/kg PO, SID for 7 days), and its appetite began to improve the following day. The frequency of respiratory signs decreased, with complete resolution of wheezing and coughing on Day 12 and 16, respectively. The duration of respiratory signs in the other symptomatic animals was shorter, lasting only 1-5 days (Table 1).

One lion (L2) developed gastrointestinal (GI) signs 10 days after the resolution of its respiratory signs. The animal refused to eat and vomited a small amount of its food. Treatment with maropitant (160 mg tablets; Zoetis Inc.; 0.8 mg/kg PO, SID for 3 days) offered in small amounts of food was initiated the following day. The animal was compliant with medication and was maintained on a reduced diet to minimize further GI upset. The lion continued to intermittently vomit frothy bile for two more days, after which time its appetite improved. The amount of food was slowly increased over the next 9 days until the lion had returned to a normal diet. No further GI signs were observed. Approximately one month after resolution of respiratory signs, one tiger (T2) vomited once and had 2 episodes of slightly loose stool over the course of a week, which then resolved spontaneously.

The tracheal wash and oropharyngeal and nasal swabs from T1 were polymerase chain reaction (PCR) negative for feline respiratory pathogens (*Bordetella, Chlamydia*, influenza, *Mycoplasma cynos, M. felis*, pneumovirus, and *Streptococcus zooepidemicus*). Cytologic examination of a direct smear and cytospin smears prepared from the flocculent tracheal wash fluid revealed epithelial necrosis and mild mixed inflammation (Fig. 3a). No infectious organisms were identified. The primary differential diagnosis for the severe necrotizing airway disease was a viral infection.

**Figure 3.**
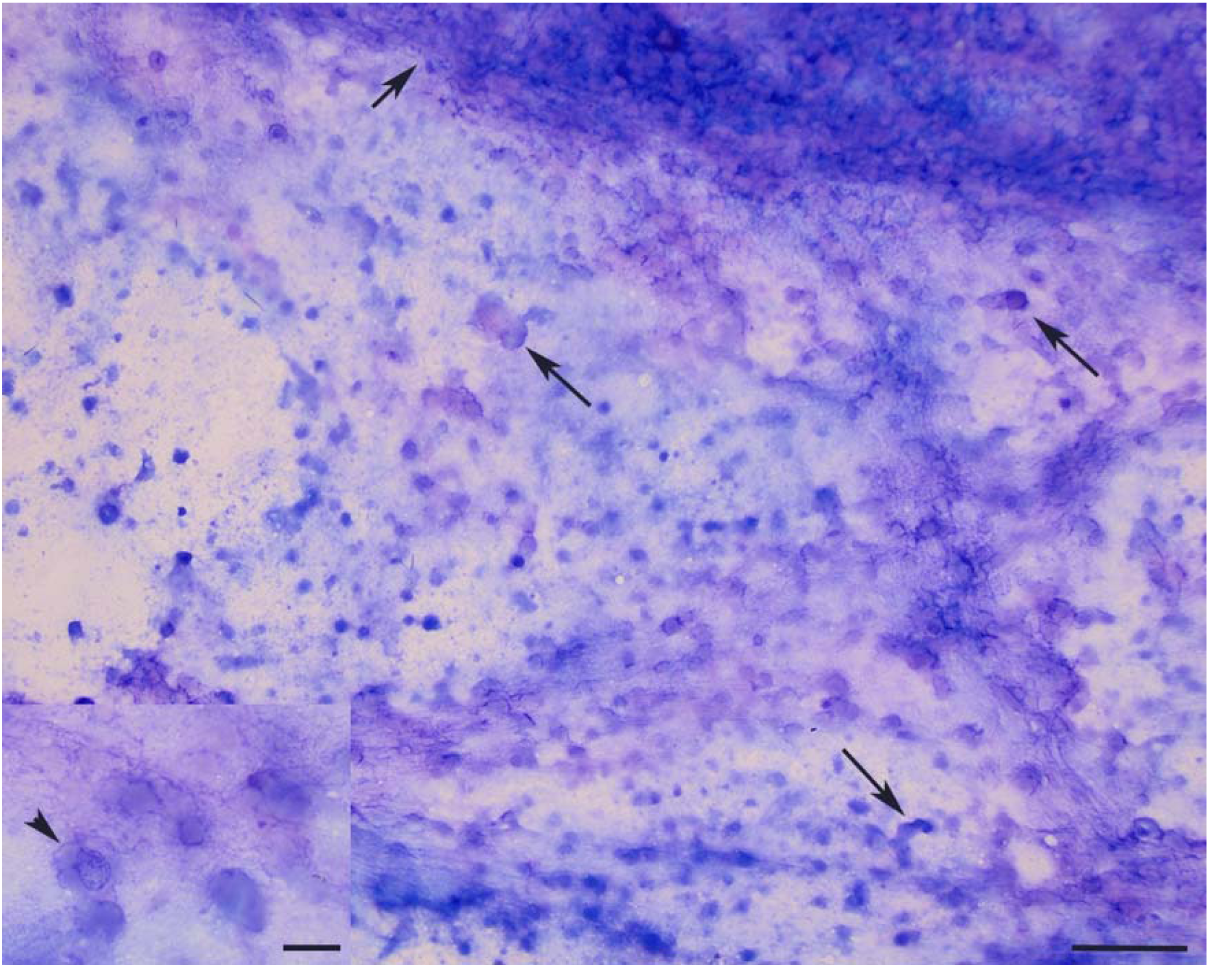
Cytologic analysis of smears from tracheal wash fluid in a tiger (*Panthera tigris jacksoni*) with SARS-CoV-2 infection. The smear contained strings of mucus (short arrow) with many enmeshed fragments and entire dying cells, several of which had a distinct columnar appearance (long arrows) with low numbers of inflammatory cells, consisting of non-degenerate neutrophils, macrophages and small lymphocytes (not shown) (modified Wright’s stain, bar = 50 um). The inset shows an epithelial cell with a faded nucleus (arrowhead) and adjacent necrotic cells or cell fragments that lack nuclei (modified Wright’s stain, bar = 12.5 um)

Specific details of initial diagnostics and sample assay methodology have been previously reported. ^12^ In brief, real-time, reverse transcription PCR (rRT-PCR) for SARS-CoV-2 performed on the tracheal wash and oropharyngeal and nasal swabs on April 3^rd^, the day after collection, yielded “presumptive positive” results for all three samples. SARS-CoV-2 infection was confirmed by rRT-PCR and partial gene sequencing on April 4 at the National Veterinary Services Laboratory.^12^ The positive result was reported to the World Organisation for Animal Health (OIE).^14^ The viral genome (designated as SARS-CoV-2/tiger/NY/040420/2020) was sequenced and aligned with SARS-CoV-2 sequences obtained from humans in NYC.^12,28^ SARS-CoV-2 was isolated from the tracheal wash sample with subsequent positive rRT-PCR results and immunofluorescent (IFA) staining for viral antigen in infected Vero cells.^12,28^ SARS-CoV-2 *in situ* hybridization (RNAScope®) was positive in a direct smear of the tracheal wash and infected Vero cells.^12^ A virus neutralization assay performed on the Day 6 serum sample from T1 yielded a titer of 1:64, confirming an immune response to infection.^12^

Daily fecal collection from the five tigers in Tiger Mountain and three lions in African Plains began on April 4, whereas daily fecal collection from the three tigers in Wild Asia began April 19. Samples from each felid were divided and frozen at – 80 °C, then sent in weekly batches to the University of Illinois Veterinary Diagnostic Laboratory (UIUC-VDL) and Cornell University’s Animal Health Diagnostic Center (AHDC) for rRT-PCR, as previously described.^12^ The UIUC-VDL targeted the N2 segment of the nucleocapsid gene, while AHDC targeted the N1 and N2 segments of the nucelocapsid gene. The rRT-PCR results were confirmed by NVSL on at least one fecal sample from each animal and indicated that all the felids in Tiger Mountain and African Plains passed SARS-CoV-2 RNA in their feces, including the one asymptomatic tiger (T5). Results were negative for the three tigers in Wild Asia for the duration of testing from April 19 to May 14. For the felids that tested positive, duration of fecal viral RNA shedding varied greatly by individual (Fig. 4). The index case shed viral RNA for 14 days, including 5 days beyond the cessation of clinical signs. In contrast, the asymptomatic tiger T5 shed viral RNA for only 5 days. The tiger (T3) with the longest duration of viral fecal shedding (24 days) was asymptomatic during this time. (Fecal collection on this animal started the day after clinical signs ceased on April 3^rd^.) Interestingly, this animal had the lowest cycle threshold (Ct) values of any cat tested, indicating the highest level of viral RNA shedding. Fecal shedding of SARS-CoV-2 RNA was also prolonged in two lions (L1 and L2) persisting for more than 30 days. L1 had marked fluctuation in shedding; it was positive on most samples collected for the first 16 days, then tested PCR negative for 13 days, then shed moderate levels of viral RNA again for two days. In L2, shedding was documented throughout the duration of this animal’s episode of GI upset (April 14 to 16). Virus was isolated from feces of two animals: from T3 on April 8^th^ (5 days after clinical signs ceased) and from L3 on April 4^th^ (the last day of clinical signs in that animal). This finding indicated that feces contained potentially infectious virus and not just viral RNA.^12^ Genome sequencing of the viral RNA in the feces showed that the tigers and lions were infected by different SARS-CoV-2 genotypes, suggesting they were infected in unrelated transmission events.^12^

**Figure 4.**
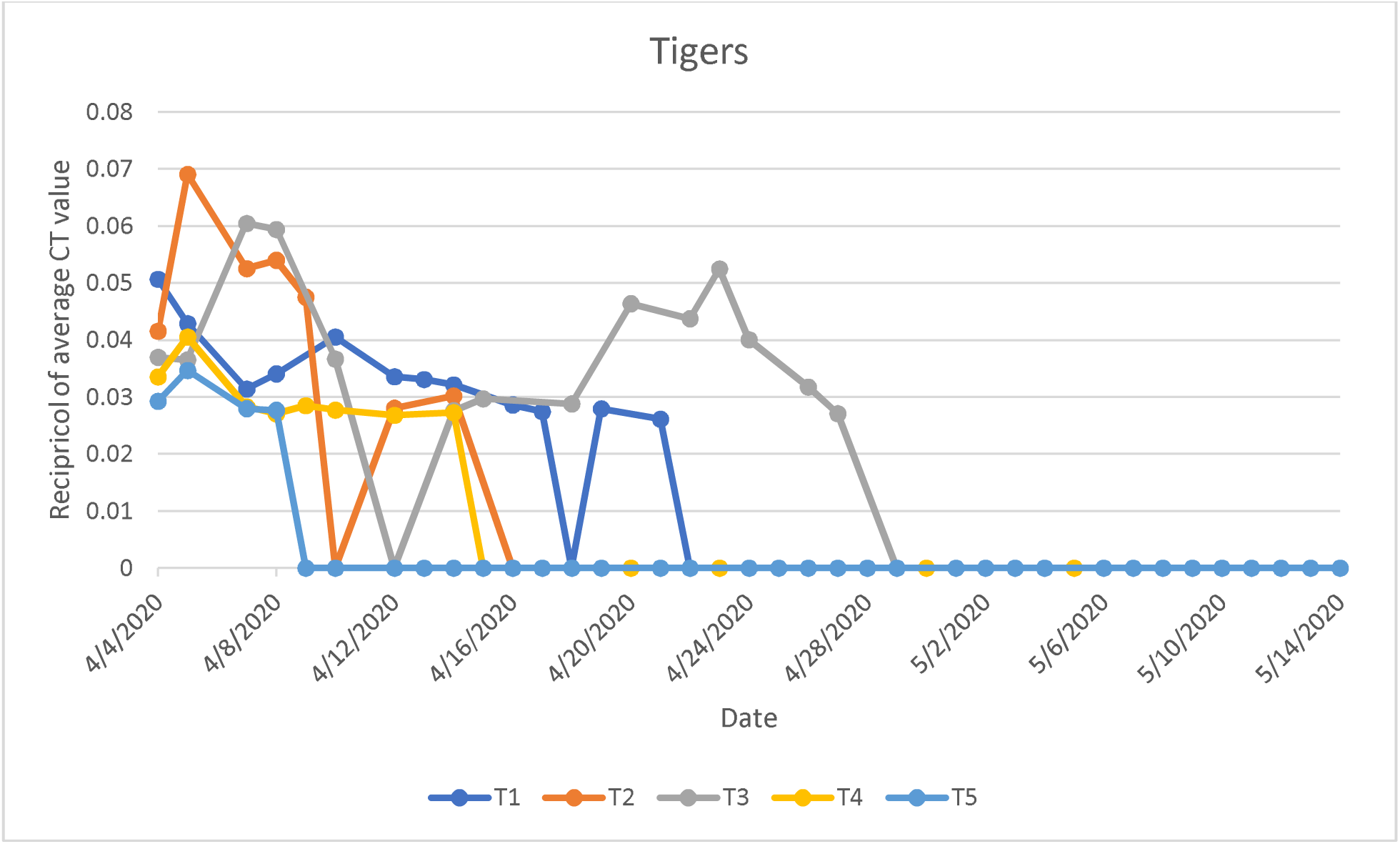
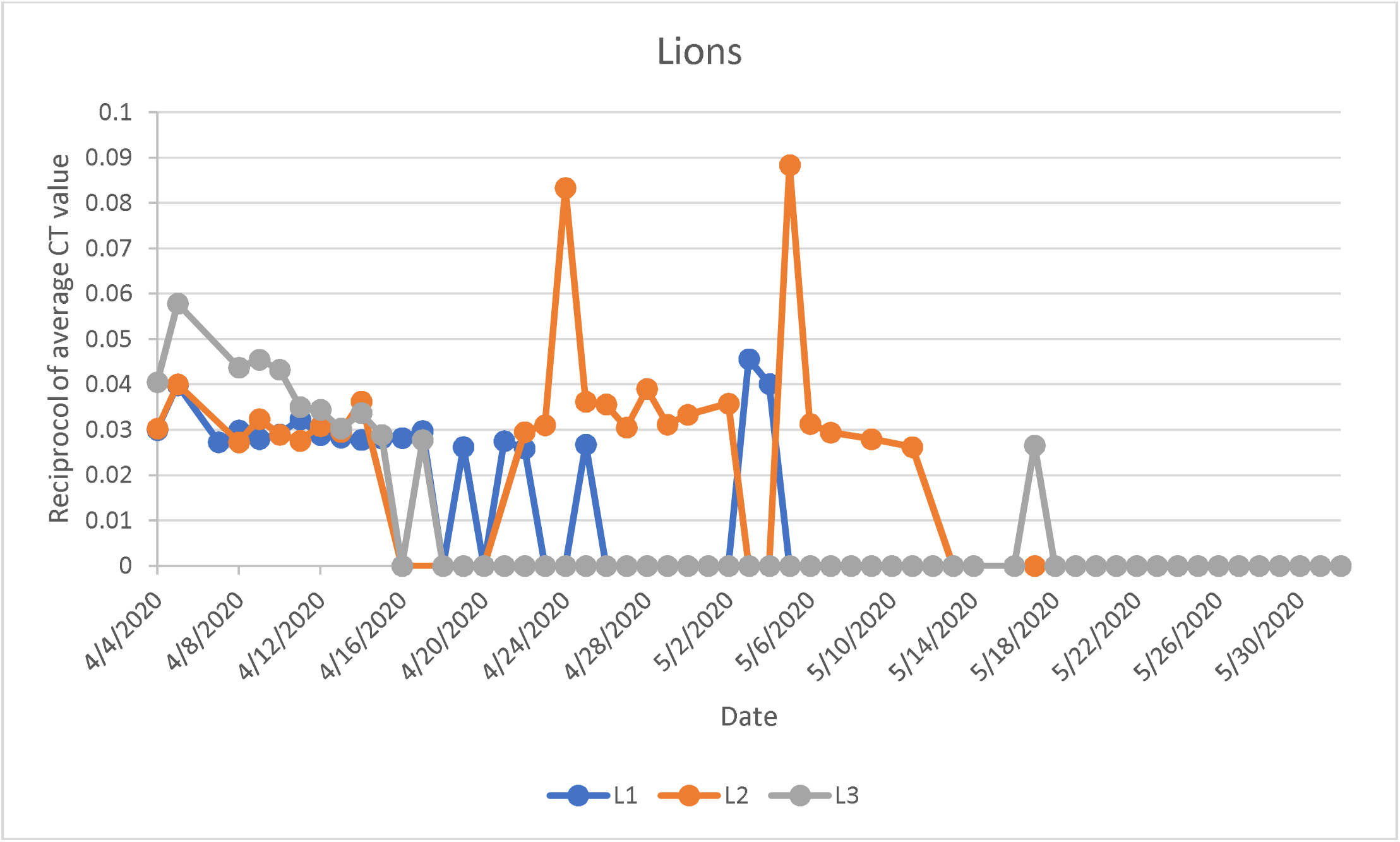
Longitudinal fecal SARS-CoV-2 RNA shedding in tigers (A) and lions (B). rRT-PCR targeting the nucleocapsid gene (segments N1 and N2 at Cornell University’s Animal Health Diagnostic Center; N2 at University of Illinois Veterinary Diagnostic Laboratory) was performed on fecal samples collected daily. The y-axis represents the reciprocal of the average cycle threshold value for all N gene segments. The positive result on L3 on May 17 was interpreted as a false positive.

Testing of keepers that worked closely with the animals in Tiger Mountain and African Plains revealed that two keepers in Tiger Mountain were PCR positive for the same strain of SARS-CoV-2 that was detected in the tigers.^12^ Two keepers who worked with the lions in African Plains were negative for viral RNA but had SARS-CoV-2 antibodies. Due to the lack of viral RNA in these keepers, the relatedness of the strain infecting the keepers and the lions could not be determined.^12^ No keepers with clinical signs of COVID-19 reported to work, in compliance with organizational policies instituted during the pandemic; transmission likely occurred during times keepers were asymptomatically shedding virus.

Upon confirmation of SARS-CoV-2 infection in T1, enhanced PPE protocols that exceeded those recommended by the AZA Felid Taxon Advisory Group (TAG) were implemented for staff working in close proximity to all felids at WCS.^38^ By the time tigers and lions at the Bronx Zoo developed clinical respiratory signs, a suspected natural SARS-CoV-2 infection in a domestic cat (*Felis catus*) had been reported in Europe.^20^ In an abundance of caution and to minimize the potential risk of additional human to felid or possible felid to human infection, staff members utilized surgical face masks, disposable gloves, eye protection (goggles or face shield), and dedicated coveralls for routine management and husbandry procedures in felid facilities at all WCS zoos including diet preparation, feeding, and cleaning. Management strategies were also altered to minimize potential exposure including limiting staff member access to the felids, implementing social distancing from the felids including keeping a minimum distance of 6 feet during shifting and feeding where possible, and temporarily discontinuing training activities.

Additional safety methods were instituted, including dry cleaning enclosures whenever possible. When hosing was required, the area was dry cleaned first, then Rescue^®^ peroxide cleaner (Virox Animal Health, Oakville Ontario L6H 6R1, Canada, 1:64 dilution in water), effective for SARS-CoV-2 disinfection, was applied to the area and allowed 5 minutes of contact time, per the manufacturer’s recommendation. Gentle hosing was then done, with avoidance of high pressure hosing to minimize aerosolization of fecal, urinary, or other waste materials.

Elective veterinary procedures and diagnostic sample collections were postponed in felids until the pandemic abated, with procedures only being performed when deemed medically necessary. During these procedures veterinary staff wore disposable gloves and surgical masks at a minimum for blood sample collection, and eye protection and N95 face masks for performing endotracheal intubations. Veterinary technicians processing fecal, blood, and urine samples in the laboratory wore gloves, masks, and eye protection.

## DISCUSSION

This report describes the clinical outcomes and SARS-CoV-2 fecal shedding patterns and duration in naturally infected tigers and lions. Detection of viral RNA by rRT-PCR and *in situ* hybridization, and of infectious virus by viral isolation in the tracheal wash confirmed the suspicion of SARS-CoV-2 infection in a tiger (T1), which was based on clinical signs of respiratory disease and cytological evidence of epithelial necrosis in the tracheal wash smears. The radiographic bronchial changes seen in the caudal thorax support an active respiratory infection. Combined with the ultrasonographic evidence of lung consolidation and coalescence of vertical artifacts (B-lines) into a “white lung” appearance, all of these changes are consistent with imaging results described for human COVID-19 patients.^23,35^ The nature of the clinical signs and presence of columnar epithelial cells in the tracheal wash smears suggested upper respiratory involvement (e.g. tracheitis or bronchitis), as well as the lower respiratory involvement documented by the imaging studies. Tracheal epithelial necrosis and pneumonitis have been reported in domestic cats experimentally infected with SARS-CoV-2.^21^

Infection was presumed for an additional three tigers in Tiger Mountain and three lions in African Plains based on temporally associated clinical respiratory signs that were similar to those of the index case (T1), fecal viral RNA shedding, and successful virus isolation from the feces of T3 and L3. Infection was not definitively confirmed in all the felids since it was elected, in the interest of human and animal safety, not to immobilize these animals solely for the purpose of collecting respiratory tract samples for testing. It is possible that the viral RNA detected represented exposure without infection in some of these felids. However, the similar clinical signs, isolation of infectious virus from feces, and moderate to large amounts of viral RNA in feces supports active infection. In addition, viral RNA was still shed in feces, particularly in T3 (which had the highest fecal load of viral RNA) long after viral RNA was no longer detected in feces from the index tiger, T1. One tiger (T5) did not demonstrate overt clinical signs of respiratory disease, yet SARS-CoV-2 RNA was detected in the feces. This animal, along with T2, did have a brief episode of epistaxis. It is unknown if this was related to coronavirus infection or other infectious agents, or to other possible causes such as trauma or dry warm ambient temperatures irritating the nasal passages. In domestic cats inoculated intranasally with SARS-CoV-2, virus was detected in the nasal turbinates within 3-6 days of inoculation, however, epistaxis was not reported in these cats.^21^ Two domestic cats in New York State were documented to have naturally acquired SARS-CoV-2 infections from suspected human to animal transmission approximately two weeks after the tiger infection was confirmed. These cats demonstrated mild respiratory illness. There was no mention of epistaxis or other clinical signs.^26^

The duration of SARS-CoV-2 RNA shedding in feces in these non-domestic felids, with T3, L1 and L2 shedding for more than 3 weeks, was noteworthy, considering that domestic cats experimentally inoculated with SARS-CoV-2 ceased shedding within a week.^21^ In humans infected with SARS-CoV-2, RNA fecal shedding frequently persists beyond 3 weeks, and shedding persists after cessation of clinical signs or viral RNA detection in sputum, similar to that seen in these non-domestic felids.^29,34^ For T3 the amount of detected viral RNA over the course of 24 days did not fit with a scenario of swallowing expectorated virus, in which progressively tapering viral RNA would be expected after cessation of clinical signs. Instead viral shedding fluctuated in this animal, with progressively decreasing amounts of viral RNA detected in feces collected from April 7 to 18, then increasing viral RNA in feces from April 19 to 22. These data support active replication of virus in the intestinal tract versus ingestion of virus from respiratory secretions. Similarly, the increase in viral RNA in the feces in L1 suggests intestinal replication, although the fluctuations in viral load in both animals may be secondary to sampling or testing variations. When L1 began shedding viral RNA again in the feces after testing negative for 13 consecutive days, neither L2 nor L3 was shedding viral RNA at the time.

It is not known if the episodes of GI clinical signs in L2 and T2 were associated with coronavirus infection or another condition. Gastrointestinal signs have been noted in a small number of human SARS-CoV-2 cases as well as in a suspected case in a domestic cat in Belgium.^1,9,20^ The GI signs occurred 10 days after coughing had ceased in L2, however the lion was still shedding SARS-CoV-2 RNA in the feces during this time. Tiger 3 had stopped shedding viral RNA in the feces several weeks prior to its episode of vomiting and soft stools. Infection with SARS-CoV-2 presumably caused the decreased appetite in T1.

The initial infection route for the tigers and lions appears to be via different keepers who were shedding virus, either due to an asymptomatic infection or before developing symptoms. Prior to T1’s onset of illness, staff members were never in shared spaces with the tigers and lions, although they did have close contact with them as part of routine training, enrichment, management, and husbandry procedures. Tigers T1 and T2 were hand raised and particularly interactive with the staff, which may have increased the likelihood of direct infection by zoo keepers. At that time, there was a recognized low risk of disease transmission between humans and felids (domestic or non-domestic), and standard practice across veterinary, curatorial, keeper and other disciplines did not recommend wearing face masks while servicing tigers, lions or other non-domestic felids. The tigers in Wild Asia, which did not show any signs of infection and consistently tested negative for SARS-CoV-2 by fecal PCR, were cared for by different keepers. Animal keepers that cared for the tigers in Tiger Mountain also cared for nine snow leopards in a different enclosure. The feces of one snow leopard with a chronic recurring cough tested negative for SARS-CoV-2 RNA on rRT-PCR. No other snow leopards showed clinical signs of respiratory disease and thus were not tested, however subclinical infections cannot be ruled out. It is possible that there are species-associated differences in susceptibility to infection within non-domestic felids. Infection by a zoo visitor was unlikely as the zoo had been closed for 11 days before the onset of clinical illness in T1, after which only essential zoo staff were on site. In addition, the design of the animal exhibits ensures that the public is separated from these species by more than 6 feet.

Once a lion was infected by close contact with a keeper, direct transmission between lions was likely since they were alternately housed together in pairs. Direct transmission between animals was unlikely for tigers because they are solitary by nature and were housed alone, however transmission through fomites or aerosol cannot be excluded. The virus was transmitted by aerosols from experimentally infected domestic cats to naïve cats housed in close proximity to but not in direct contact with the infected cats.^21^ None of the tigers were ever in the same enclosure at the same time. T1 was housed adjacent to T2, and they alternated access to dens and a common yard. Tigers T3, T4, and T5 did not share common areas with T1 and T2, which could suggest a common source of infection from a keeper, or transmission by aerosol or fomites. T4 was housed in the middle of the building, adjacent to T1/T2 and T3/T5 dens on either side. However, T3, T4, and T5 were shifted through common spaces in order to access a common outdoor yard. Tigers were not allowed access to the exhibit from the onset of clinical signs in T1 through May 1 so that the animals could be more closely monitored. Fomite or surface contact transmission between these tigers is possible. Other potential sources of infection were food contamination during diet preparation before PPE was implemented and infectious aerosols generated by felid vocalizations or cleaning procedures.

Although there were no confirmed or peer reviewed reports of natural SARS-CoV-2 infections in other zoo or wildlife species at the time of this documented infection at the Bronx Zoo, there was concern about the potential susceptibility of other taxa due to a report of experimental inoculation of domestic cats and ferrets with SARS-CoV-2.^21^ Due to these concerns, and in accordance with the guidelines recommended by the AZA Great Ape TAG veterinary advisors, staff members at WCS continued to adhere to previously established internal PPE guidelines for primates, including wearing surgical masks and gloves when servicing all primates, and wearing eye protection when working with Old World primates including western lowland gorillas (*Gorilla gorilla gorilla*).^37^ Disposable gloves and masks were also used for food preparation. Gloves and surgical mask use was implemented for servicing and food preparation for small carnivores of the Orders Viverridae, Herpestidae, Mustelidae, and Mephitidae, as well as Chiroptera, as recommended by the AZA Small Carnivore and Bat TAGs.^39,40^ Elective procedures and diagnostic sampling were suspended in these taxa and only medically necessary procedures continued.

The decision to expand the use of PPE in non-human primates, bats, and small carnivores was based in part upon previous documentation of infections of SARS-CoV-1 and SARS-CoV-like viruses in some of these taxa. *Rhinolophus* bat species are natural reservoirs for SARS-like viruses.^10^ SARS-CoV-like viruses have been isolated from or viral RNA detected in Himalayan palm civets (*Paguma larvata*) and raccoon dogs (*Nyctereutes procyonoides*); the virus sequence was 99.8% similar to the SARS-CoV-1 that caused a human epidemic in 2002-2003.^27^ Civets were also experimentally infected with two different SARS-CoV-1 isolates.^5,32^ Chinese ferret badgers (*Melogale moschata*) produced neutralizing antibodies after natural exposure to SARS CoV-1.^27^ Experimental infection with SARS-CoV-1 was demonstrated in ferrets (*Mustela furo*), cynomolgus macaques (*Macaca fascicularis*), rhesus macaques (*Macaca mulatta*), and African green monkeys (*Chlorocebus aethiops*).^3,11,19,24^ Non-peer reviewed publications also reported SARS-CoV-2 RNA shedding in two naturally exposed dogs in Hong Kong.^20^ Natural infection of SARS-CoV-2 with associated respiratory signs was also reported in farmed mink in the Netherlands and Denmark.^17,18^

To date, no other non-domestic felids or other animals at the Bronx Zoo or any of the other WCS zoos and aquarium have become ill due to a confirmed SARS-CoV-2 infection. Although there were anecdotal reports of other felids at zoological institutions in the US and abroad that had possible clinical signs of SARS-CoV-2 infections, to our knowledge there has been only one other confirmed infection in a non-domestic felid: a puma at a zoo in South Africa.^15^ It is unknown why with the high numbers of captive non-domestic felids and the high prevalence of COVID-19 infections in humans throughout much of the world, two independent transmission events from humans to non-domestic cats would occur at one location in one week but rarely elsewhere.

## CONCLUSIONS

This case series confirms susceptibility of tigers and lions to SARS-CoV-2. Clinical signs include coughing, wheezing, and inappetence, and possibly vomiting and epistaxis. The course of disease in tigers and lions at the Bronx Zoo was generally short, with coughing usually resolving within 5 days but in one case continuing for 16 days. SARS-CoV-2 RNA was detected by rRT-PCR in oropharyngeal and nasal swabs, tracheal wash fluid, and feces in the index tiger, and virus was isolated from tracheal wash fluid in the index tiger and feces from another tiger and lion.^12^ Fecal viral RNA shedding persisted for as long as 35 days beyond cessation of respiratory signs, suggesting viral replication in the GI tract. Asymptomatic infection was suspected in one tiger via the detection of SARS-CoV-2 RNA in feces. Fecal testing has the advantage of being non-invasive, and can be an effective way to screen animals. The index tiger also demonstrated seroconversion on a virus neutralization test. Serologic testing would be useful for screening non-domestic felids for SARS-CoV-2 exposure and infection, and such testing is planned for the lions and tigers in this case series. However, testing methods currently rely on virus neutralization, which requires biosafety level-3 conditions in the laboratory, and other serologic methods need to be developed and validated for such testing to be rigorously conducted. No additional tigers, lions or other non-domestic felids at any of the WCS zoos developed clinical signs after implementation of new PPE protocols despite ongoing high levels of human infection and community spread in NYC through June 2020. Personal protective equipment should therefore be used as a means to minimize the chance of anthropozoonotic transmission of coronaviruses to Felidae and other susceptible taxa. SARS-CoV-2 is an OIE reportable disease, and current recommendations for testing animal samples include coordination with state regulatory agencies.

## Acknowledgements

The authors are truly grateful for the dedication and expertise of the staff at the Wildlife Conservation Society’s Bronx Zoo Department of Mammalogy including Ralph Aversa, Mary Gentile, Michelle Medina, Phil Reiser, Brent Atkinson, Jennifer Cott, Lauren DelGrosso, David Fernandez, Kristin Nielsen, Chris Salemi and Amanda Scherer; and Zoological Health Program, including Jessica Long and Dr. Jean Pare; and the diagnostic teams at UIUC-VDL and AHDC labs and NVSL. Special thanks also goes to the state animal and public health officials in New York, Drs. Smith, Newman and Slavinski, and Illinois, Drs. Ernst and Austin, for facilitating rapid actions.

